# LSD1 represses a neonatal/reparative gene program in adult intestinal epithelium

**DOI:** 10.1101/2020.02.20.958363

**Authors:** Rosalie T. Zwiggelaar, Håvard T. Lindholm, Madeleine Fosslie, Marianne T. Pedersen, Yuki Ohta, Alberto Díez-Sánchez, Mara Martín-Alonso, Jenny Ostrop, Mami Matano, Naveen Parmar, Emilie Kvaløy, Roos R. Spanjers, Kamran Nazmi, Morten Rye, Finn Drabløs, Cheryl Arrowsmith, John Arne Dahl, Kim B. Jensen, Toshiro Sato, Menno J. Oudhoff

**Affiliations:** CEMIR – Centre of Molecular Inflammation Research, Department of Clinical and Molecular Medicine, NTNU – Norwegian University of Science and Technology, 7491 Trondheim, Norway; Department of Microbiology, Oslo University Hospital, Rikshospitalet, NO-0027, Oslo, Norway; BRIC: Biotech Research and Innovation Centre, University of Copenhagen, DK-2200 Copenhagen N, Denmark; Department of Gastroenterology, Keio University School of Medicine, Tokyo 160-8582, Japan; Department of Organoid Medicine, Keio University School of Medicine, Tokyo 160-8582, Japan; Department of Oral Biochemistry, Academic Centre for Dentistry (ACTA), 1081LA Amsterdam, The Netherlands; Department of Clinical and Molecular Medicine, NTNU – Norwegian University of Science and Technology, 7491 Trondheim, Norway; Clinic of Surgery, St. Olav’s Hospital, Trondheim University Hospital, 7030 Trondheim, Norway; Structural Genomics Consortium, University of Toronto, Toronto, ON, M5G 1L7, Canada; Princess Margaret Cancer Centre, University Health Network, Toronto, ON, M5G 2M9, Canada; Department of Medical Biophysics, University of Toronto, Toronto, ON, M5G 1L7, Canada; Novo Nordisk Foundation Center for Stem Cell Research, Faculty of Medical and Health, University of Copenhagen, DK-2200 Copenhagen N, Denmark

## Abstract

Intestinal epithelial homeostasis is maintained by adult intestinal stem cells, which, alongside Paneth cells, appear after birth in the neonatal period. We aimed to identify new regulators of neonatal intestinal epithelial development by testing a small library of epigenetic modifier inhibitors in Paneth cell-skewed organoid cultures. We found that Lysine-specific demethylase 1A (*Kdm1a/Lsd1*) is absolutely required for Paneth cell differentiation. *Lsd1*-deficient crypts, devoid of Paneth cells, are still able to form organoids without a requirement of exogenous or endogenous Wnt. Mechanistically, we find that LSD1 represses genes that are normally expressed in fetal and neonatal epithelium. This gene profile is similar to what is seen in repairing epithelium, and indeed, we find that *Lsd1*-deficient epithelium has superior regenerative capacities after irradiation injury. In summary, we found an important regulator of neonatal intestinal development and identified a druggable target to reprogram intestinal epithelium towards a reparative state.

## INTRODUCTION

The intestinal epithelium undergoes a dramatic change during the neonatal period. Crypt formation occurs after birth together with the appearance of Paneth cells (PCs) and the development of adult intestinal stem cells (ISCs). Adult ISCs rely on niche factors such as Wnt ligands. *In vivo*, mesenchymal cells are important sources of Wnt to support ISC maintenance (Degirmenci et al., 2018; Greicius et al., 2018; Shoshkes-Carmel et al., 2018), whereas *in vitro*, it is PCs that are required to supply the necessary Wnt (Sato et al., 2011; Durand et al., 2012; Kim et al., 2012; Farin et al., 2012). In contrast, Wnt ligands or PCs are dispensable for fetal organoid homeostasis (Fordham et al., 2013; Mustata et al., 2013). Thus, ISCs undergo a fetal-to-adult transition that includes a change in Wnt dependency.

Recently, the existence of bona fide fetal ISCs has been challenged by the finding that any fetal epithelial cell can be or become an adult ISC as long as the appropriate environment is supplied (Guiu et al., 2019). This model fits nicely with studies showing that after injury, the intestinal epithelium is temporarily reprogrammed into a fetal-like state that is needed for proper repair (Yui et al., 2018; Nusse et al., 2018; Gregorieff et al., 2015). This, in turn, complements work specifying that adult intestinal epithelial lineages can dedifferentiate to give rise to new ISCs to rebuild the epithelium after injury (Tetteh et al., 2016; Yu et al., 2018; van Es et al., 2012; Buczacki et al., 2013). In hindsight, these high levels of cell-fate reversion make sense because the intestine is a common site for chemical and mechanical challenges as well as the host for many putative pathogens. Nonetheless, it is not yet fully understood how the fetal-to-adult ISC transition, or its reversal upon injury, is mediated, and whether epithelial reprogramming can be targeted therapeutically.

Adult ISCs give rise to all intestinal epithelial subtypes. Unlike other stem cell systems, both ISCs and differentiating intestinal epithelial cells have a similar chromatin state at lineage defining genes, which allows for Notch-mediated lateral inhibition in ISC differentiation (Kim et al., 2014). However, the same group subsequently identified differences in open chromatin distinguishing ISCs from secretory precursors, which was reversed upon irradiation when these secretory precursors dedifferentiate into ISCs (Jadhav et al., 2017). Three groups separately identified a crucial role for the polycomb repressive complex 2 (PRC2) in maintaining crypt physiology (Jadhav et al., 2019; Koppens et al., 2016; Chiacchiera et al., 2016). In summary, although it is clear there is an important role for certain epigenetic modifiers in intestinal epithelial biology, the role and importance of others remain undefined. Here we exploit the availability of chemical probes targeting epigenetic modifiers (Scheer et al., 2018), and combine it with intestinal organoid cultures (Sato et al., 2009), to identify the demethylase LSD1 as a critical regulator of crypt maturation including PC differentiation.

## RESULTS AND DISCUSSION

### Identification of LSD1 as a regulator of Paneth cells

The intestinal epithelium undergoes a dramatic change during the neonatal period including the appearance of Paneth cells (PCs). To study PC differentiation in organoids, we developed a differentiation protocol (adapted from (Yin et al., 2013)) using CHIR (GSK3 inhibitor) and DAPT (γ-secretase inhibitor) to activate Wnt and block Notch signalling, respectively (Fig. S1A). CHIR-DAPT treatment led to a robust enrichment of Lysozyme^+^ PCs by confocal microscopy, mRNA, and protein expression compared to standard EGF, Noggin, and R-spondin 1 (ENR) organoid growing conditions (Fig. 1A, S1B). Next, we tested chemical probes targeting 18 methyltransferases and demethylases and identified the LSD1 inhibitor GSK-LSD1 to consistently repress PC differentiation (Fig. 1B, 1C, S1C, S1D, Supplementary table 1). In support, GSK-LSD1 similarly affected PC differentiation in organoids grown in ENR conditions (Fig. 1D, S1E), and use of a different LSD1 inhibitor led to a similar near loss of PCs (Fig. S1F). Consistent with the irreversible binding nature of GSK-LSD1 to its target (Mohammad et al., 2015), we found that upon withdrawal of GSK-LSD1, PCs re-appeared after 2 organoid splitting events (Fig. 1E, S1G).

**Fig. 1.**
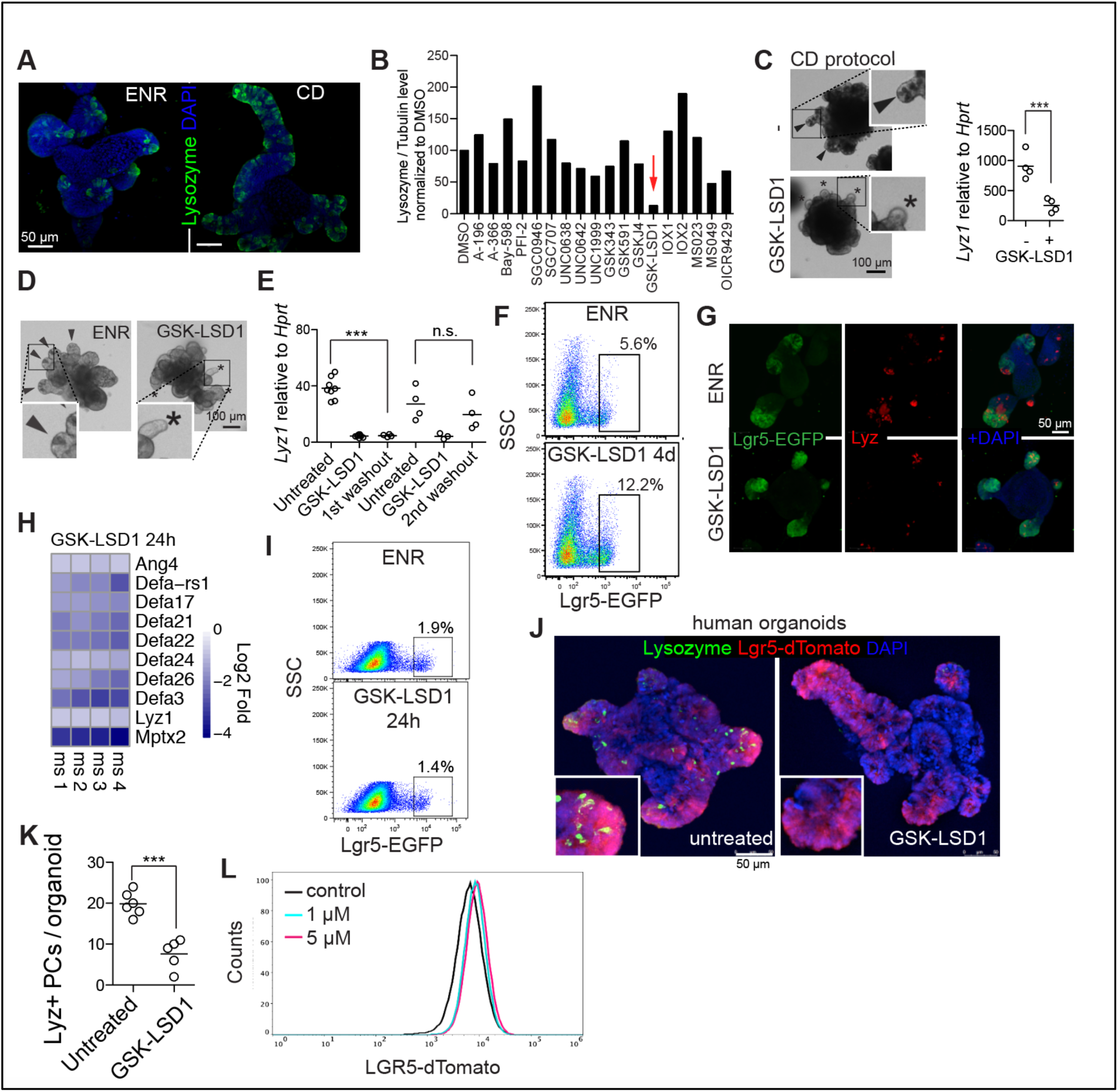
Inhibition of LSD1 blocks PC differentiation and allows niche-independent expansion of ISCs. **A**, Confocal images of Lysozyme and DAPI staining of ENR and CHIR-DAPT (CD protocol as in (S1A)) grown organoids. **B**, Quantification of Lysozyme/Tubulin levels by western blot of organoids treated with indicated inhibitors in CD conditions. Concentrations used can be found in supplementary table 1. Data is mean from 2 independent experiments (see Fig. S1C, D for raw data). **C**, Brightfield images and inserts of CD-grown organoids with and without GSK-LSD1 (1 µM). *Lyz1* expression relative to *Hprt* by qPCR. **C&D**, Arrows indicate PC^+^ crypts based on presence of granular cells, asterisks indicate PC^-^ crypts. **D**, Brightfield images and inserts of ENR-grown organoids with or without GSK-LSD1 (1 µM). **E**, *Lyz1* expression relative to *Hprt* by qPCR. **F & I**, Flow cytometry of Lgr5-EGFP organoids. Representative plot from at least n=3 different mouse lines. **G**, Images of Lgr5-EGFP (anti-GFP) and Lyz (Lysozyme) staining of organoids. **H**, Heatmaps of indicated genes from RNA-seq of 24 h GSK-LSD1 treated organoids. N=4 different mouse organoid lines, and expression is relative to each individual control. **J&K**, Images of human LGR5-dTomato organoids, additionally stained for Lysozyme. Quantified number of PCs per human organoid from 2 different experiments. **L**, Flow cytometry of LGR5-dTomato treated human organoids with 1 or 5 µM. ** p<0.01 ***p<0.001

### Inhibition of LSD1 renders Lgr5^+^ cells independent of niche-providing PCs *in vitro*

PCs are crucial for adult small intestinal organoids as they supply niche factors to retain a stem cell population, and normally, PC-devoid organoids only sustain growth upon Wnt supplementation (Sato et al., 2011; Durand et al., 2012; Kim et al., 2012; Farin et al., 2012). To test the role of LSD1 in intestinal stem cells, we used Lgr5-EGFP derived organoids and treated them with GSK-LSD1. We found that GSK-LSD1 treatment resulted in a 2-3 fold increase in percentage of Lgr5^+^ cells (Fig. 1F, 1G). Thus, GSK-LSD1 treatment expands the Lgr5^+^ population and renders these cells independent of PC-derived niche factors such as *Wnt3* (Fig. S1H). To test if only a selection or all PC genes are downregulated upon LSD1 inhibition, we performed RNA-seq on organoids treated with GSK-LSD1 for 24 h, between day 1 and 2 after splitting, which we anticipate is right when PCs develop. We found a robust downregulation of all PC specific genes (gene set from (Haber et al., 2017)) (Fig. 1H). In contrast, 24 h GSK-LSD1 treatment did not lead to an expansion of the Lgr5^**+**^ population (Fig. 1I). Thus, the expansion of Lgr5^**+**^ cells upon LSD1 inhibition does not precede or outcompete PC loss and may be a separate event. Nevertheless, using a recently described culture condition that allows PC differentiation in human organoids (Fujii et al., 2018), we found that GSK-LSD1 also blocks PC differentiation in human organoids while simultaneously expanding the LGR5^**+**^ population (Fig. 1J-L).

### LSD1 is required for PCs but not goblet or enteroendocrine cells *in vivo*

To test the role of LSD1 *in vivo*, we crossed *Lsd1*^f/f^ (Kerenyi et al., 2013) with *Villin-*Cre (Marjou et al., 2004) mice to delete *Lsd1* in intestinal epithelial cells specifically (KO mice). Although these mice appear normal, we found that KO mice lack PCs throughout the small intestine (Fig. 2A, S1I). We did observe that KO mice had ‘escaper’ crypts still expressing LSD1, and LSD1^+^ crypts were positive for Lysozyme (Fig. S2J). Thus, these mice do not completely lack PCs. Currently, two genes are known to be absolutely required for PC differentiation *in vivo; Sox9* and *Atoh1* (Bastide et al., 2007; Yang et al., 2001). We did not observe differences in Sox9 expression (Fig. 2A), and, although we found fewer *Atoh1*^+^ cells in KO epithelium (Fig. 2A), reduction of *Atoh1*^+^ cells unlikely causes a complete lack of PCs. We reasoned that perhaps KO crypts are filled with PC precursors expressing *Wnt3*, however, similar to GSK-LSD1 treated organoids, *Wnt3* was markedly reduced in crypts of KO mice crypts (Fig. 2A). Next, we examined other intestinal secretory lineages and found a reduction of goblet cells, but equal numbers of enteroendocrine cells, comparing adult WT and KO littermates (Fig. 2B). When we examined fetal and postnatal intestines of WT and KO littermates, we found that the reduction in goblet cells emerges after developmental stage P7, similar to when PCs start to develop (Fig. 2C, 2D, S2K). These results suggest that LSD1 KO epithelium maintains neonatal characteristics into adulthood, including the absence of PCs and fewer goblet cells.

**Fig. 2.**
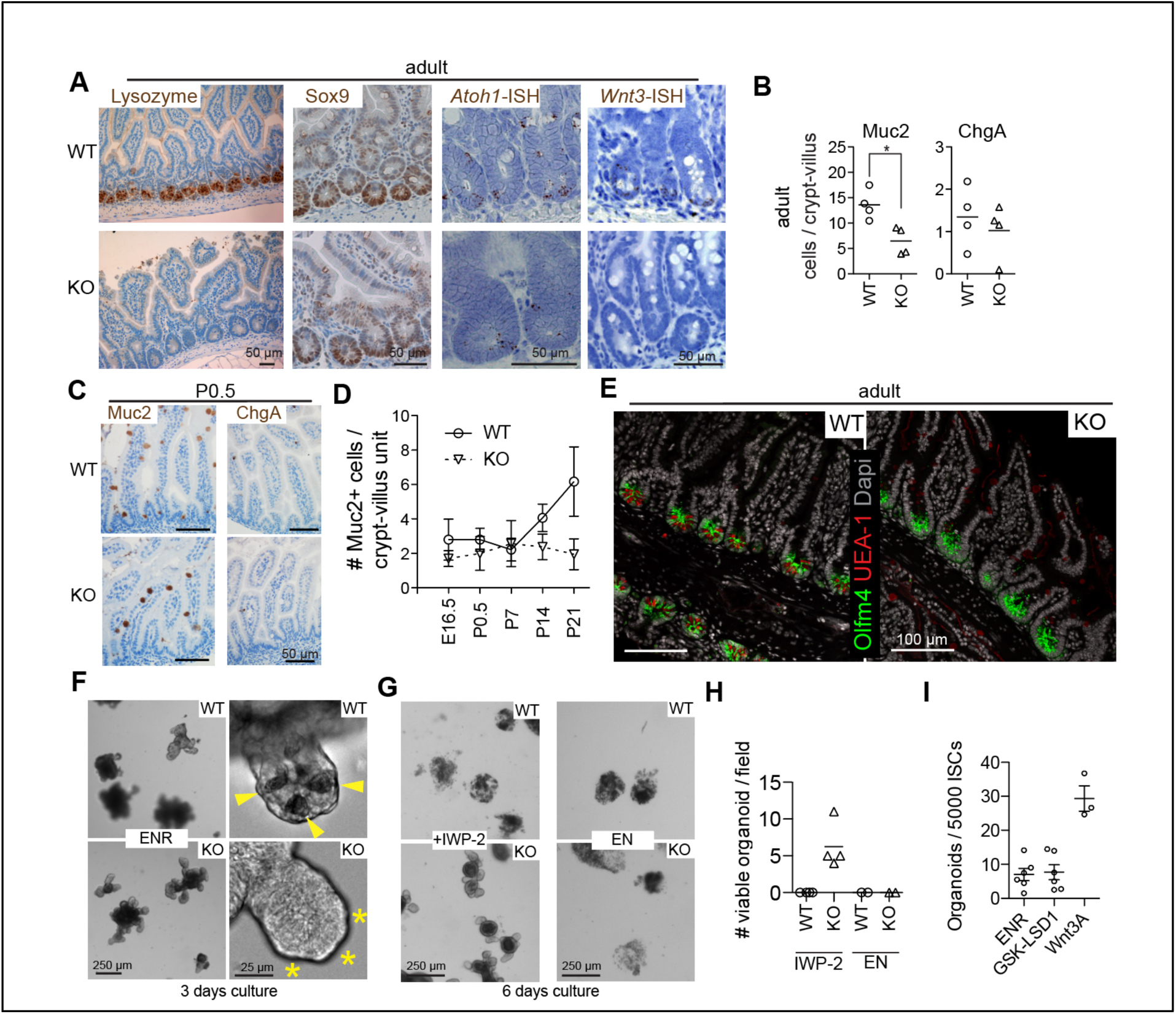
LSD1 is required for crypt maturation *in vivo* and Wnt dependency of organoids. **A**, Representative images of antibody (Lysozyme and Sox9) and *in situ* hybridization (ISH) (*Atoh1* and *Wnt3*) staining of WT and KO small intestinal tissue. **B**, Quantification of Muc2^**+**^ goblet and ChgA^**+**^ enteroendocrine cells in adult duodenum intestine. N=4 mice, *p<0.05. **C**, Representative images of Muc2 and ChgA antibody staining at P0.5 **D**, Quantifications of Muc2^**+**^ goblet cells throughout development. N ≥ 3 mice, mean ± SEM is shown. **E**, Representative image of Olfm4 antibody and UEA-1 staining of adult WT and KO tissue. **F-H**, Brightfield images of WT and KO organoids with indicated treatments, quantified are wells from 2 different experiments (n=2 mice). **I**, Organoid outgrowth from single sorted Lgr5^HIGH^ ISCs, each dot represents a mouse, data pooled from 3 independent experiments. Mean and SE are shown.

### Mice lacking LSD1 sustain crypt-bottom ISCs independent of PCs, and KO organoids grow independent of endogenous Wnt

To test the role of LSD1 in ISCs *in vivo*, we used *in situ* hybridization (ISH) and antibody-based detection of the ISC marker *Olfm4* in tissues of WT and KO mice. We found Olfm4^+^ cells completely filling the bottom of crypts in KO mice, compared to the standard PC/ISC pattern observed in WT crypts (Fig. 2E, S1L). In addition, all crypt-base cells in KO mice are Ki67^+^, suggesting that these Olfm4^+^ cells are proliferating (Fig. S2L). *Atoh1*^-/-^ mice lack PCs, and *Atoh1*^-/-^ crypts do not sustain organoid growth without Wnt supplementation (Durand et al., 2012). In contrast, *Lsd1* WT and KO crypts were equally able to form organoids, even in the absence of PCs in KO organoids (Fig. 2F). This led us to hypothesize that KO organoids do not rely on endogenous Wnt. Indeed, blockage of Wnt signaling by the porcupine inhibitor IWP-2, showed that treated KO organoids sustained growth unlike WT organoids (Fig. 2G, 2H). Of note, IWP-2 distinctively reduced growth rate in KO organoids, which makes long-term expansion unfeasible, yet, after splitting there were still surviving KO organoids under continuous IWP-2 treatment, and, LSD1 inhibitor treatment greatly increased splitting efficiency (Fig. S1I, S1J). In contrast, both KO and WT organoids could not sustain growth in medium lacking R-spondin 1 (Fig. 2G, 2H). Thus, loss of LSD1 activity in ISCs renders them not requiring (endogenous) Wnt for growth and even leads to expansion of this population (Fig. 1). However, GSK-LSD1 is not able to replace Wnt3A in the ability to form organoids from single ISCs (Fig. 1I). Although KO organoids resemble aspects of those derived from fetal epithelium that also lack PCs (Fordham et al., 2013; Mustata et al., 2013), there are some key distinctions. Unlike fetal organoids, KO organoids cannot grow without R-spondin 1. In addition, KO organoids form crypts, whereas fetal or Wnt-supplemented organoids mainly grow as spheroids.

Our data thus suggests that adult KO epithelium is in between fetal and adult: KO organoids do not rely on Wnt yet are unable to grow without R-spondin 1, and KO epithelium *in vivo* lack PCs and have reduced GC numbers yet with crypts that have Olfm4^**+**^ ISCs.

### LSD1 represses fetal and neonatal genes that allows PC differentiation independent of YAP/TAZ

Next, we sought to find the mechanism by which LSD1 controls intestinal epithelial biology. We performed RNA-seq on FACS-sorted Epcam^**+**^ small intestinal crypt cells from WT and KO mice. We found 2564 up and 1522 down regulated genes (p-adj<0.1) in KO cells compared to WT cells (Fig. 3A). In support of our findings that there are *Atoh1*^+^ cells in KO crypts and the differential ability of *Atoh1*-KO and *Lsd1-*KO crypts to form organoids, we found no shift towards an *Atoh1*^-/-^ gene signature in the KO transcriptional profile (Fig. S2A) (Kim et al., 2014). To verify our gene expression profile, we tested a PC-specific gene signature (Haber et al., 2017), and, expectedly, found that this is repressed in KO crypts (Fig. 3B).

**Fig. 3.**
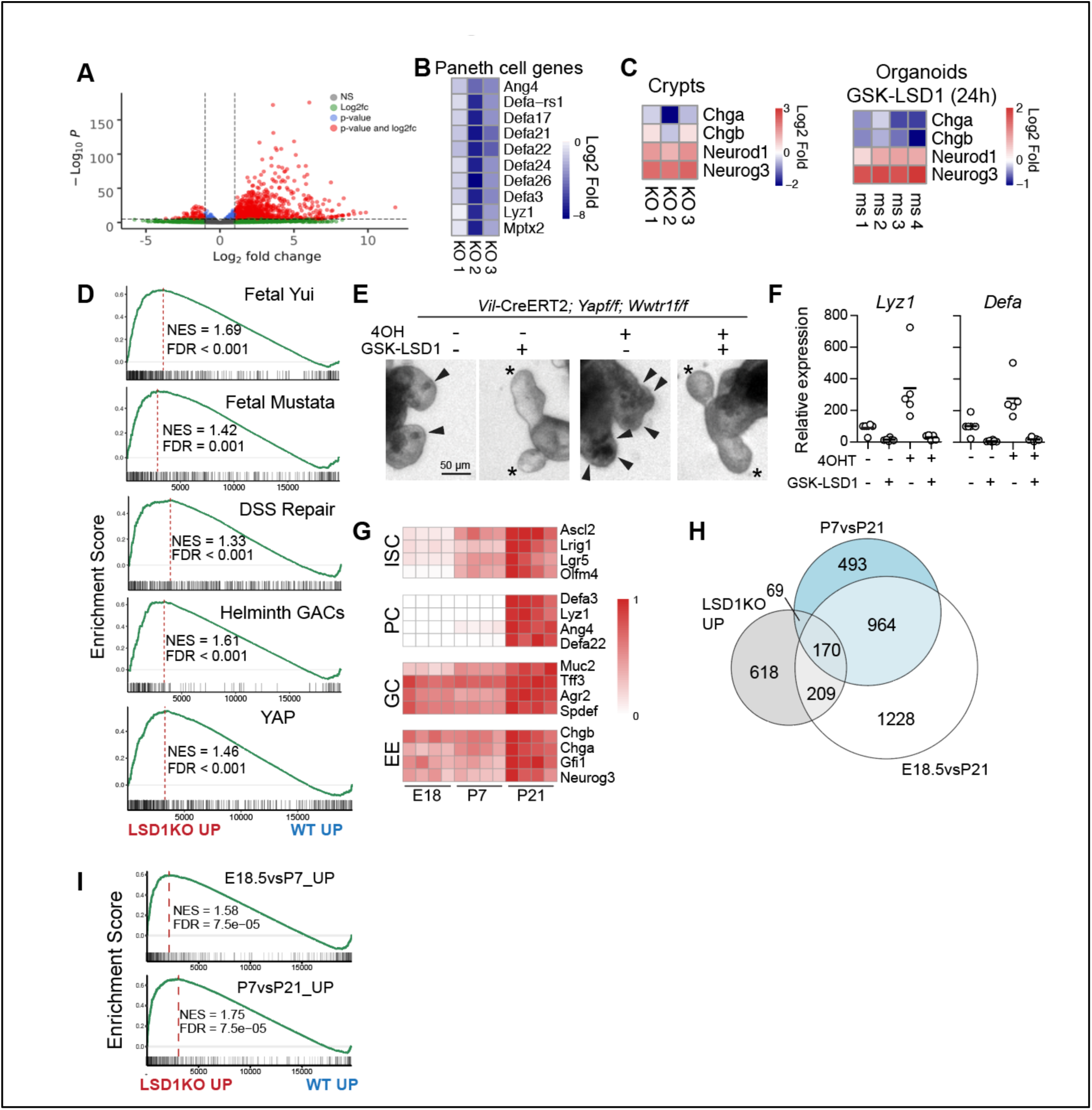
Deletion of LSD1 renders intestinal epithelium fetal-like. **A**, Volcano plot of RNA-seq data comparing WT and KO crypt cells (n=3). **B**, Heatmap of PC-specific genes. **C**, Heatmaps of enteroendocrine associated genes from indicated RNA seq data. **D & I**, LSD1KO *vs.* WT RNA-seq data was analyzed by gene set enrichment analysis (GSEA) for indicated gene signatures. Normalized Enrichment Score (NES) and False Discovery Rates (FDR) are indicated. **E**, Brightfield images of organoids with indicated treatments. Arrows indicate granular PCs, asterisks indicate crypts devoid of PCs. **F**, Expression of *Lyz1* and *Defa* by qPCR relative to *Actb*, each dot represents a well, pooled from 4 independent experiments. **G**, Heatmap of genes associated with ISCs, PCs, goblet cells (GC), and enteroendocrine cells (EE) during development. Shows expression relative to the highest TPM for that gene, which is set to 1. **H**, Venn diagram of genes significantly higher expressed in LSD1KO compared with genes that are up in P7*vs*.P21 and E18*vs.*P21.

A study reported that an LSD1-GFI1 complex was rapidly disturbed by LSD1 inhibitors, and that the scaffolding role of LSD1 rather than its enzyme activity was most crucial (Maiques-Diaz et al., 2018). Notably, *Gfi1*^-/-^ mice have a near loss of PCs and ISCs, reduced goblet cells, but more enteroendocrine cells (Shroyer et al., 2005; Sato et al., 2011). When we examined genes known to be repressed by GFI1 in intestinal epithelium, we indeed found rapid and robust increase of *Neurog3* and *Neurod1*, but not that of mature enteroendocrine cell markers *Chga* and *Chgb* (Fig. 3C). This is in accordance with equal enteroendocrine cells *in vivo* in WT and KO mice (Fig. 2C, S1K). Together, this suggests that inhibition of LSD1 rapidly leads to enzyme-independent de-repression of GFI1-targeted enteroendocrine-progenitor markers, that in turn can limit PC differentiation (Shroyer et al., 2005). Furthermore, these enteroendocrine progenitors may take over the role of PCs in Notch-dependent crypt formation (van Es et al., 2019). However, it does not explain the expansion of ISCs, the complete loss of PCs, or the lack of expansion of mature ChgA^**+**^ cells that we observe in KO epithelium, which thus contradict a primarily GFI1-mediated mechanism.

Over the years, there have been various different intestinal stem cell and stem-cell like populations described, either using genetic markers such as Lgr5 or Bmi1, or by techniques such as label-retaining capacity and scRNA-seq (Yan et al., 2017; Buczacki et al., 2013; van Es et al., 2012; Ayyaz et al., 2019; Muñoz et al., 2012). To test if there was enrichment for a certain stem cell type, we analyzed expression of defining genes for each population (Fig. S2B, S2C), however, we did not find enrichment for any stem cell subtype including the classical Lgr5^**+**^ ISC signature. Instead, we find enrichment of two fetal gene sets in the KO transcriptional profile (Fig. 3D) (Yui et al., 2018; Mustata et al., 2013). Recently, two groups elegantly identified and characterized a cellular repair state in the intestinal epithelium that is fetal-like (Yui et al., 2018; Nusse et al., 2018). Indeed, two repair gene signatures from these studies were also enriched in the KO transcriptional profile (Fig. 3D). Yui *et al.* revealed that the reparative state was mediated by YAP/TAZ (Yui et al., 2018). A separate YAP gene signature was enriched in our KO transcriptional profile (Fig. 3D) (Gregorieff et al., 2015).

To test if YAP/TAZ is required for LSD1-mediated PC differentiation, we treated organoids derived from mice lacking both *Yap* and the gene encoding for TAZ *Wwtr1* (*Vil-* Cre;*Yap*^f/f^;*Wwtr1*^f/f^) with GSK-LSD1 and found that PC differentiation was equally impaired in WT and mutant organoids (Fig. S2D). However, we noted that approximately half of these organoids contained *Yap* and *Wwtr1* based on qPCR (Fig. S2E). We thus tested a second model, deleting YAP/TAZ in an inducible manner (*Vil-*CreERT2; *Yap*^f/f^;*Wwtr1*^f/f^ organoids). Indeed, tamoxifen treatment led to near undetectable levels of both *Yap* and *Wwtr1* (Fig. S2F), impaired survival and led to an increase in PCs (Fig. 3E, 3F), in accordance with previous results (Azzolin et al., 2014; Gregorieff et al., 2015). Strikingly, GSK-LSD1 treatment completely abrogated PC differentiation independently of YAP/TAZ (Fig. 3E, 3F).

To distinguish fetal from postnatal intestinal epithelial development we performed RNA-seq on embryonic (E) day 18.5, as well as postnatal (P) days 7 and P21 intestinal epithelium (Fig. S2G, S2H). Figure 3G shows how established cell lineage markers behave during development. As expected, we observed a stepwise increase in ISC markers, an abrupt appearance of PC genes at P21, and goblet and enteroendocrine gene expression at E18.5 and P7 stages that only increases slightly at P21 (Fig. 3G). This indeed supports our hypothesis that KO mice have intestinal epithelium that is ‘stuck’ at a P7 stage, lacking PCs, with reduced goblet cell and immature enteroendocrine cells, but with crypts containing ISC-like cells. When we compared genes upregulated in KO crypts to different developmental stages we see overlap with both E18.5 and P7 stages compared to P21 (Fig. 3H). In a separate test, we found enrichment by GSEA for our own fetal (E18vsP7) and neonatal (P7vsP21) gene sets (Fig. 3I).

### LSD1 controls H3K4me1 levels of genes associated with a fetal-like profile

LSD1 controls embryonic development by repressing enhancers to allow for embryonic stem cell differentiation (Agarwal et al., 2017; Whyte et al., 2012). We did not observe major differences in global H3K4me1/2 levels by immuno-histochemistry comparing escaper LSD1^+^ crypts in KO tissue (Fig. S3A). To assess if LSD1 mediates H3K4 demethylation, we performed ChIP-seq for H3K4me1 and H3K4me2 comparing sorted WT and KO crypt cells. Analysis of ChIP-seq for H3K4me1 identified 2059 sites with associated altered methylation levels, of which the majority (1622) were enriched in KO crypts (Fig. 4A). ChIP-seq of H3K4me2 revealed a very similar pattern and the large majority of genes affected by LSD1 with gain of H3K4me2 were also significant in the analysis for H3K4me1 (Fig. 3A, 3B). In addition, most of these peaks are not in close proximity to TSS sites (Fig. S3B). The top 300 genes associated with increased H3K4me1 levels in the KO, are overall enriched in the KO transcriptional profile (Fig. 4C). Furthermore, using a high stringency (adj. p<0.01), we combined our RNA-seq and H3K4me1 ChIP-seq data to establish a core list of 228 genes that are mediated by the enzymatic activity of LSD1 (Table S2). Importantly, 84% of the increased H3K4me1 peaks associated with these 228 genes are located outside the 2 kb surrounding the TSS. This suggests that LSD1 would drive enhancer-mediated regulation of these genes, which fits with a role generally associated with LSD1 (Whyte et al., 2012; Agarwal et al., 2017). Importantly, the LSD1 core signature is enriched in a transcriptional profile comparing fetal with adult organoids (Fig. 4D (Yui et al., 2018)). Thus, we propose that LSD1 enzymatically represses genes that are required for maturation of intestinal epithelium.

**Fig. 4.**
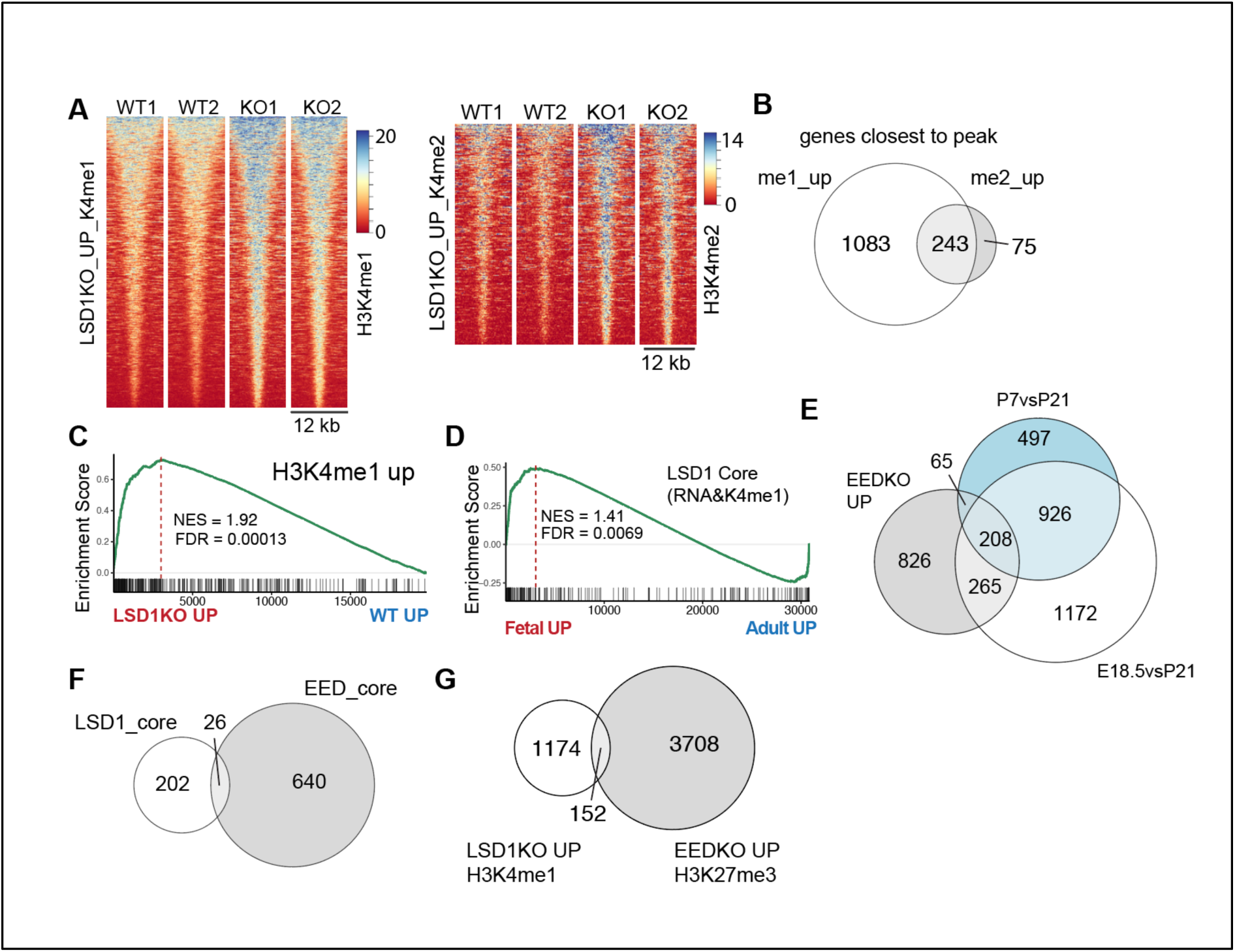
LSD1 controls H3K4me1/2 levels of fetal-like genes. **A & C**, Heatmaps of H3K4me1 and H3K4me2 sites that are significantly up in KO crypts compared to WT crypts. **B**, Venn diagram comparing genes associated with H3K4me1 and H3K4me2 peaks that were significantly higher in KO crypts compared to WT crypts. **C**, RNA-seq analysis by GSEA of KO *vs* WT transcriptional profile on a gene set consisting of genes with associated increased H3K4me1 levels in KO crypts. **D**, GSEA of LSD1 core gene list (228 genes) on a transcriptome data set that compares fetal to adult organoids (Yui et al., 2018). **E**, Venn diagram of genes significantly higher expressed in EEDKO compared with genes that are up in P7*vs*.P21 and E18*vs.*P21.**F & G**, Venn diagrams comparing the LSD1 core, and the EED core (genes up in EED KO crypts AND have H3K27me3 associated peak) as well as the genes associated with increased methylation levels in each separate group. locus at indicated developmental stages. **F**, Heatmap of H3K4me1 sites that were identified to be significantly higher at E18 (UP) compared to P21 (DOWN) (left panel). Heatmap of WT and KO H3K4me1 profiles of

### LSD1 controls genes separately from PRC2

Several groups have shown that EED, an essential component of the Polycomb Repressive Complex 2 (PRC2), is essential for maintaining adult ISCs and crypts, likely by repressing fetal and embryonic genes (Koppens et al., 2016; Jadhav et al., 2019; Chiacchiera et al., 2016). Indeed, EED controlled genes show a remarkable similar overlap with fetal and neonatal genes as LSD1 controlled genes (Fig. 4E, 3H). However, where EED KO epithelium returns to a fetal and even embryonic state and mice become moribund (Jadhav et al., 2019), LSD1 KO mice retain an early-life postnatal stage and appear normal up to at least 1 year. Fittingly, comparing the LSD1 core with the EED core (genes up in EEDKO crypts and associated with H3K27me3 peaks (Koppens et al., 2016)) revealed strikingly little overlap between regulated genes (Fig. 4F). Further analysis confirmed that also the majority of putative LSD1-controlled H3K4me1 genes are not ‘co-repressed’ by the PRC2-mediated H3K27me3 (Fig. 4G). Thus, this suggests that both LSD1 and PRC2 control fetal-like genes but in an unrelated fashion.

### LSD1 expression is downregulated during repair

We conclude that LSD1 represses fetal/neonatal genes in adult epithelium. A similar gene set is re-activated after damage during the repair phase. Thus, we hypothesized that during repair de-repression of LSD1-controlled fetal/neonatal genes would require LSD1 inactivation.

Therefore, we studied LSD1 expression after whole body irradiation (10 gy) and found that LSD1 is lower expressed in large crypts that appear 3 days post irradiation, compared to normal-sized crypts of naïve mice (Fig. 5A). Of note, this is opposite of the Hippo-transducer YAP expression pattern (Fig. S3C). In support, we assessed *Lsd1* expression in a recently described single cell RNA-seq experiment comparing crypt cells during homeostasis and during active repair (Ayyaz et al., 2019), and found a clear reduction in number of crypt cells expressing *Lsd1* (Fig. 5B). In summary, *Lsd1* is actively downregulated in the repair phase of intestinal epithelium.

**Fig. 5.**
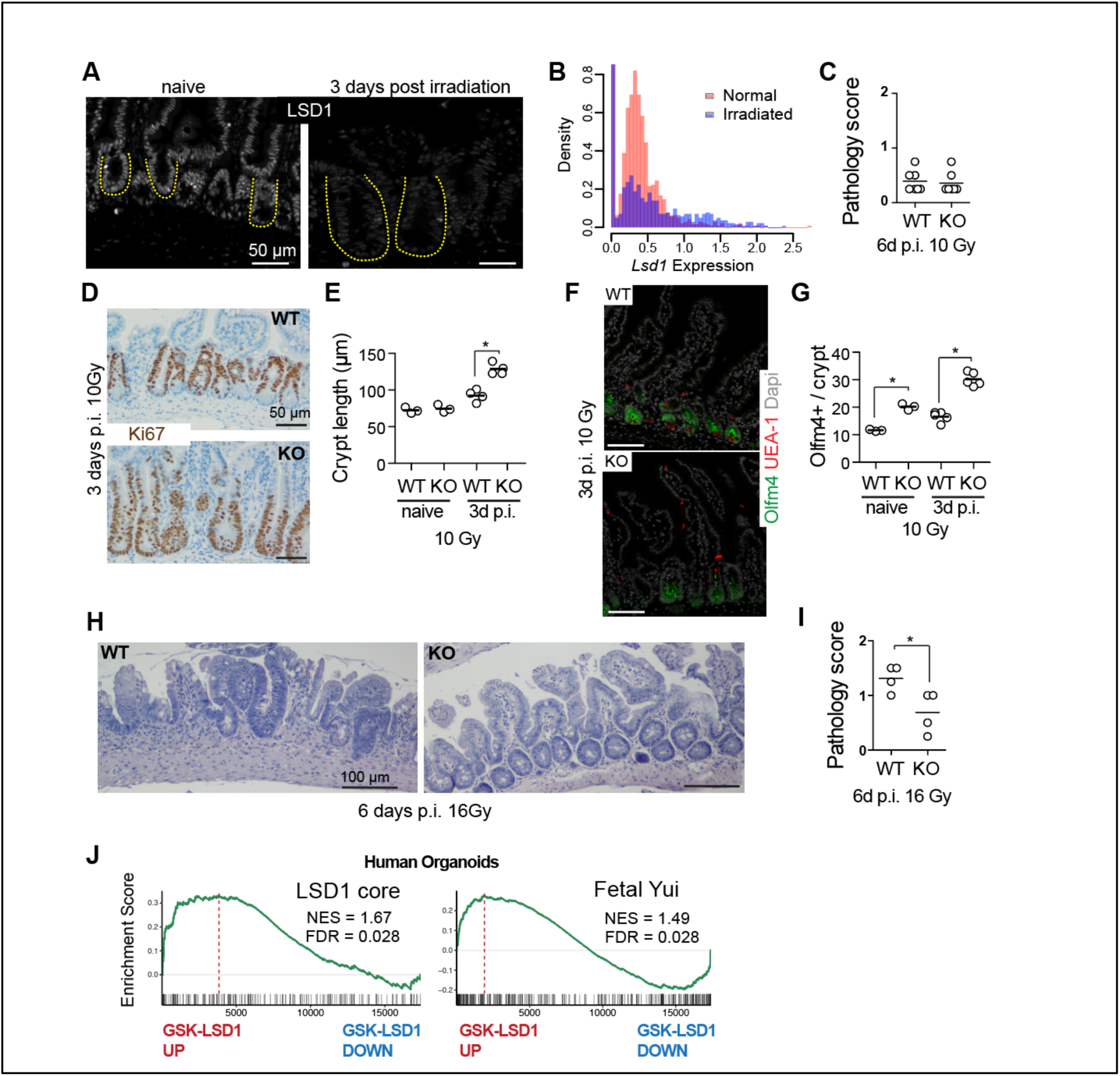
LSD1 levels are actively reduced during repair and *Lsd1*-deficient epithelium repairs better than WT tissue. **A**, Images of LSD1 antibody staining of naïve and irradiated (10 gy) intestines. **B**, *Lsd1* expression from single-cell RNA-seq data from (Ayyaz et al., 2019) in normal and irradiated crypts. **C**, Pathology score of indicated intestines 6 days post irradiation (p.i.) with 10 Gy. **D**, Images of Ki67 antibody staining of WT and KO tissue 3 days p.i.. **E**, Crypt length as quantified from images shown in 6D and S2H, crypt length is determined by Ki67^**+**^ cells. **F**, Images of Olfm4, UEA-1, and Dapi stained WT and KO intestines 3 days p.i. at 10 Gy. **G**, Quantifications of Olfm4^**+**^ cells per crypt quantified from images such as shown in 6F and 2G. **H**, H&E staining of WT and KO intestinal tissue 6 days p.i. with 16 Gy. **I**, Pathology score of indicated intestines 6 days p.i. with 16 Gy. **J**, GSEA of indicated signatures on transcriptome data generated from control and GSK-LSD1 treated human organoids. * p<0.05

### *Lsd1*-deficient epithelium has superior reparative capacity

The reparative-like features of KO epithelium prompted us to test if this would be beneficial upon injury. We irradiated WT and KO mice with 10 Gy and we did not observe pathological differences 6 days post treatment (Fig. 5C, S3D). However, we did find an increase in crypt length 3 days after injury, as measured by Ki67^+^ cells (Fig. 5D, 5E). Of note, we found no evidence of appearing PCs after injury in KO tissue (Fig. 5F), but we did observe that the Olfm4^**+**^ ISC zone similarly expanded as the Ki67^+^ crypts (Fig. 5F, 5G). This suggests that KO epithelium may be have better reparative capacities, but with this level of irradiation WT epithelium is also able to recover after 6 days. Thus, we next irradiated mice with 16 Gy, when WT mice are unable to recover by day 6, and in striking contrast, KO epithelium regained crypt-villus structures and had lower pathology scores compared to WT epithelium (Fig. 5H, 5I, S3E, S3F). Thus, KO epithelial tissue that have a pre-existing repairing profile enhances repair *in vivo* after radiation injury.

### Sustained inhibition of LSD1 renders human organoids fetal-like

Importantly, the role we find for LSD1 suppressing fetal/neonatal genes is the opposite of what recently was found in skin, where inhibition of LSD1 led to increase of fate-determining transcription factors and increase in differentiation, which was beneficial for treatment of skin cancer (Egolf et al., 2019). This is similar to what has been proposed for other cancer types, including intestinal tumors in zebrafish, where LSD1 inhibition would lead to differentiation and reduced tumor development (Maiques-Diaz et al., 2018; Rai et al., 2010). To test the translational potential of our findings, we performed a gene expression array of GSK-LSD1 treated human organoids, and found that the LSD1 core and a fetal signature were enriched in GSK-LSD1 treated human organoids (Fig. 6J).

In summary, we provide evidence that inhibition of LSD1 may be a viable target for the reprogramming of intestinal epithelium into a reparative state that is beneficial after injury, such as inflicted by radiation therapy.

## ACKNOWLEDGEMENTS

We thank Drs. Stuart Orkin, Sylvie Robine, and Stefano Piccolo for kindly sharing mouse strains. We thank Colby Zaph for the initial support of this study and critical reading of the manuscript. We thank Unni Nonstad for assistance with cell sorting. We are indebted to Anne Marthinsen for performing the irradiation after working hours, and the Department of Radiology and Nuclear Medicine (St. Olavs Hospital) for allowing the use of their instruments. We thank the imaging (CMIC) and animal care (CoMed) core facilities that assisted in this work (NTNU). The WT KO crypt RNA-seq was done by the Genomics Core Facility at NTNU, which receives funding from the Faculty of Medicine and Health Sciences and Central Norway Regional Health Authority. The ChIP sequencing was done at the Norwegian Sequencing Centre (www.sequencing.uio.no), a national technology platform hosted by the University of Oslo and supported by the “Functional Genomics” and “Infrastructure” programs of the Research Council of Norway and the Southeastern Regional Health Authorities. Funding of this work was provided by the Norwegian Research Council (Centre of Excellence grant 223255/F50, and ‘Young Research Talent’ 274760 to MJO) and the Norwegian Cancer Society (182767 to MJO). MMA is the recipient of a Marie Skłodowska-Curie IF (DLV-794391). This work was also supported by the South-Eastern Norway Regional Health Authority, Early Career Grant 2016058, and the Research Council of Norway “Young Research Talent” grant to JAD. MF is supported by the Norwegian Research Council (grant no. 275286). The SGC is a registered charity (number 1097737) that receives funds from AbbVie, Bayer Pharma AG, Boehringer Ingelheim, Canada Foundation for Innovation, Eshelman Institute for Innovation, Genome Canada through Ontario Genomics Institute [OGI-055], Innovative Medicines Initiative (EU/EFPIA) [ULTRA-DD grant no. 115766], Janssen, Merck KGaA, Darmstadt, Germany, MSD, Novartis Pharma AG, Ontario Ministry of Research, Innovation and Science (MRIS), Pfizer, São Paulo Research Foundation-FAPESP, Takeda, and Wellcome. This project also received funding from the European Union’s Horizon 2020 research and innovation programme (grant agreements INTENS 668294, MTP, KBJ). The Novo Nordisk Foundation Center for Stem Cell Biology is supported by a Novo Nordisk Foundation grant number NNF17CC0027852 (KBJ).

## Author contributions

R.T.Z., H.T.L., M.F., M.T.P., Y.O., A.D.S, M.M.A., J.O., M.M., N.P., E.K., R.R.S., and M.J.O. performed experiments. K.N. and C.A. provided essential reagents. M.R., F.D., C.A., J.A.D., K.B.J., T.S., and M.J.O. supervised and/or provided conceptual insight. M.J.O wrote a first draft with subsequent input from all authors.

## Competing interests

T.S. is an inventor on several patents related to organoid culture.

Correspondence and request for materials should be addressed to M.J.O.

## Supplementary Figures and Full Methods

**Fig. S1.**
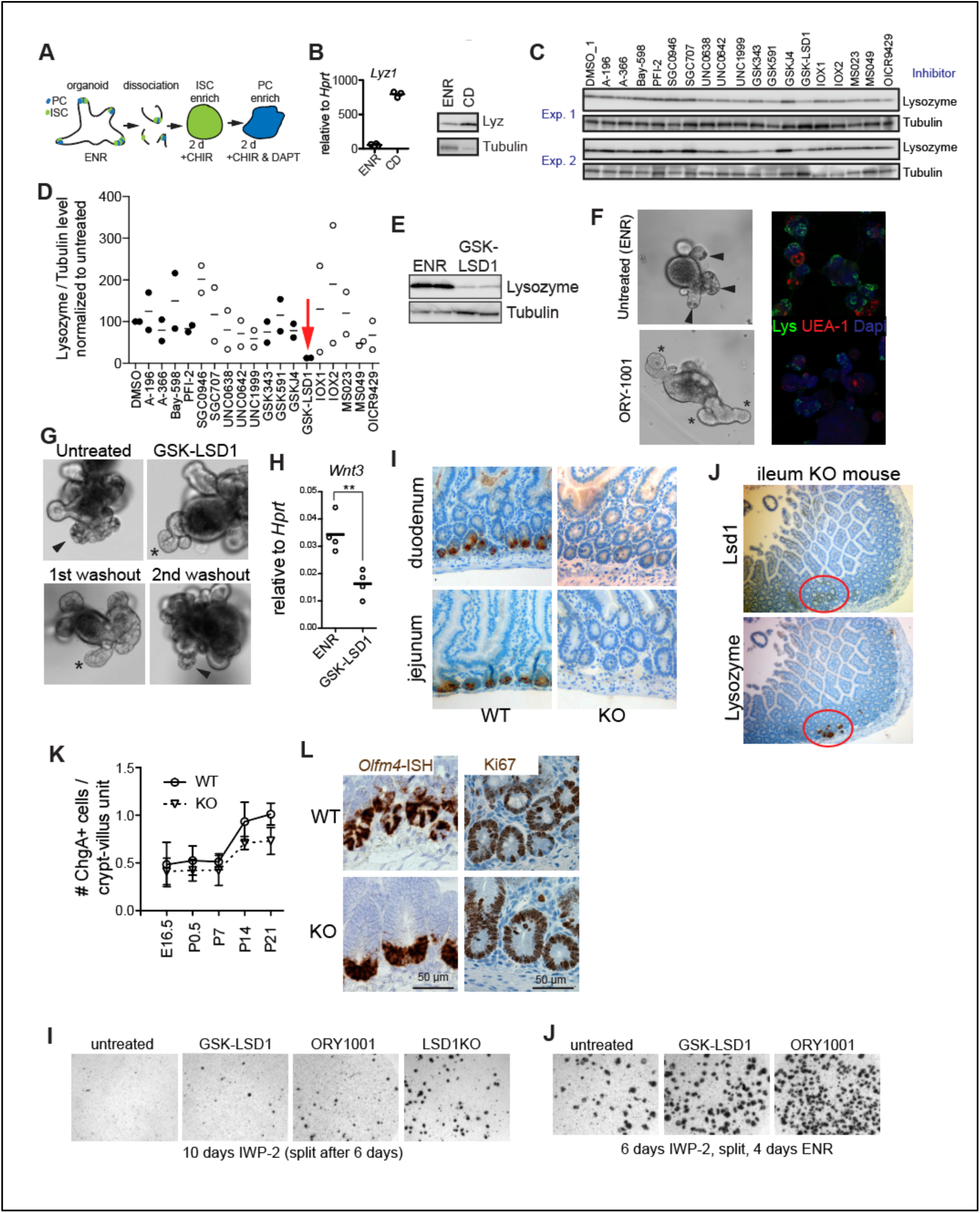
**A**, Schematic overview of organoid PC differentiation protocol. ENR (EGF, Noggin, R-spondin 1). **B**, *Lyz1* expression relative to housekeeping *Hprt* of ENR *vs.* CD cultured organoids. **B&C**, Western blot probed for anti-Lysozyme and anti-Tubulin antibodies of organoid lysates from ENR *vs.* CD cultured organoids, and CD treated organoids treated with indicated inhibitors. **D**, Quantification of Lysozyme/Tubulin levels by western blot of two independent screens. **E**, Western blot of organoids cultured in ENR with or without GSK-LSD1. **F**, Organoids were treated with the LSD1 inhibitor ORY-1001 (100 nM) for 4 days. Brightfield images and confocal staining of Lysozyme (Lys in green), UEA-1 (red) and Dapi (blue) is shown. **G**, Representative images of a washout experiment. We observed the return of PCs after the 2^nd^ washout. Arrow depict granular PCs, asterisk depict crypts devoid of organoids. **H**, *Wnt3* expression relative to *Hprt.* **I**, Images of indicated WT and KO tissue stained for PC marker Lysozyme. **J**, Images of sequential sections of KO small intestine tissue showing that LSD1^**+**^ escaper crypts are Lysozyme^**+**^. **K**, Quantifications of ChgA^**+**^ enteroendocrine cells throughout development. N ≥ 3 mice, mean ± SEM is shown. **L**, Images of *Olfm4-*ISH and Ki67 antibody stained sections of duodenum tissue from adult WT and KO mice. **I & J**, Representative images of organoid cultures treated with indicated inhibitors (GSK-LSD1 5 µM, ORY1001 100 nM, IWP-2 2µM).

**Fig. S2.**
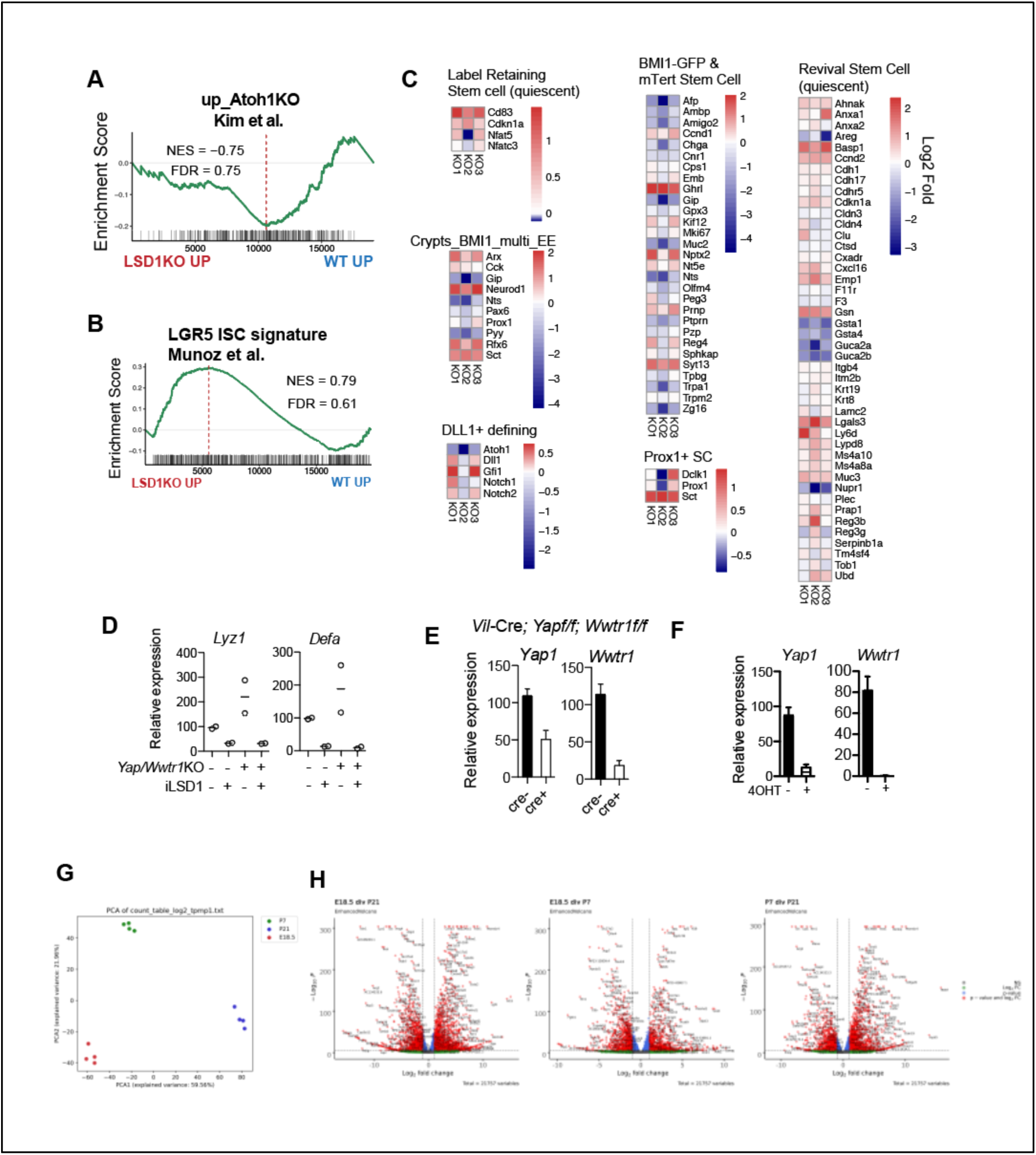
**A & B**, LSD1KO *vs.* WT RNA-seq data was analyzed by gene set enrichment analysis (GSEA) for indicated gene signature. Normalized Enrichment Score (NES) and False Discovery Rates (FDR) are indicated. **C**, Heatmaps for genes that are associated with or define different types of intestinal stem cells. **D**, qPCR of PC markers *Lyz1* and *Defa* relative to *Gapdh* of organoids from control (Cre-) or mutant (Cre+) *Vil*-Cre; *Yap1*^f/f^; *Wwtr1*^f/f^ mice treated with GSK-LSD1 (as indicated). Means from a representative experiment (n=2) is displayed, performed three times. **E**, Expression of indicated genes by qPCR relative to *Gapdh* from WT (Cre-) and KO (Cre+) organoids derived from *Vil*-Cre; *Yap1*^f/f^; *Wwtr1*^f/f^ mice. Mean + SEM from n=2. **F**, qPCR relative to *Actb* from *Vil*-CreERT2; *Yap1*^f/f^; *Wwtr1*^f/f^ treated with vehicle or 4OH-Tamoxifen (OHT) with or without GSK-LSD1, n=4. **G**, PCA plot of RNA-seq experiment comparing E18, P7, and P21 intestinal epithelium. **H**, Volcano plots of the differential expression comparing E18, P7, and P21.

**Fig. S3.**
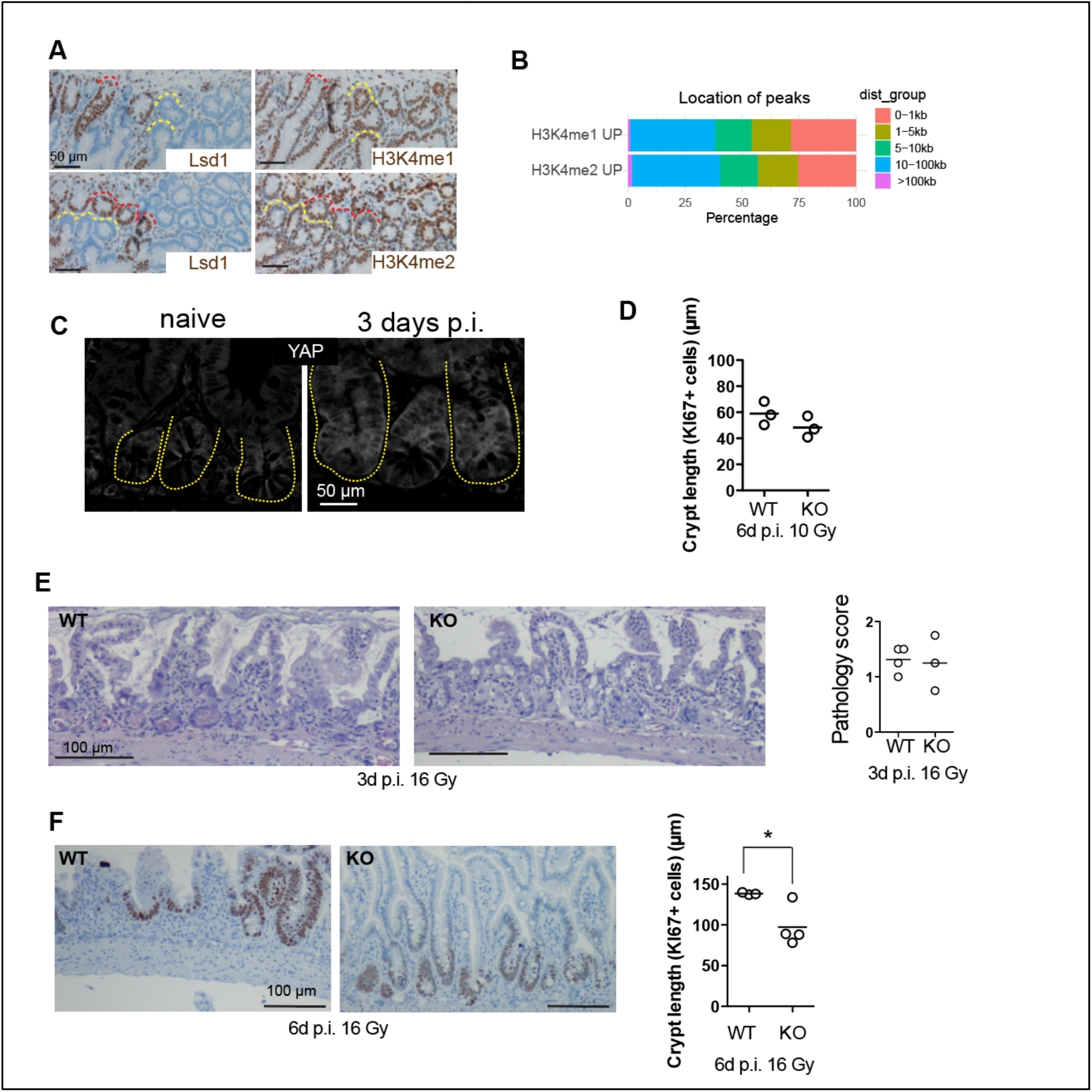
**A**, Images of immune-histochemistry of KO mice including escaper crypts. Yellow=KO crypts, Red=Escaper Lsd1^**+**^ crypts. **B**, Overview of localization relative to TSS of peaks that were up in KO crypts compared to WT crypts for indicated epitopes. **C**, YAP antibody staining of small intestinal crypts (yellow dotted lines) of untreated naive mice, and irradiated (10 gy) mice 3 days post irradiation (p.i.). **D**, Crypt length at 6 days p.i. with 10 Gy. **E**, H&E staining and pathology score of intestines 3 days p.i. with 16 Gy. **F**, Ki67 staining and quantification of Ki67+ crypts at 6 days p.i. with 16 Gy.

## METHODS

### Animal work

*Lgr5-EGFP-IRES-CreERT2* (stock no: 008875) mouse strain was obtained from Jackson Laboratories. *Villin-Cre* and *Villin-CreERT2* (Marjou et al., 2004) were a kind gift from Sylvie Robine, *Lsd1*^f/f^ mice were a kind gift from Dr. Stuart Orkin (Kerenyi et al., 2013). Mice were housed and maintained at the Comparative Medicine Core Facility (CoMed), and experiments were ethically approved by the Norwegian Food Safety Authority (FOTS ID: 11842). Mice were lethally irradiated (10 Gy or 16 Gy) and small intestinal repair was assessed 3 and 6 days post irradiation. *Yap*^f/f^;*Wwtr1*^f/f^ animals (Azzolin et al., 2014) were a kind gift from Stefano Piccolo and were crossed with *Villin-Cre* and *Villin-CreERT2* at University of Copenhagen under the approval of the National Animal Ethics Committee in Denmark.

### Crypt, IEC, and ISC isolation, mouse organoid cultures

*Adult crypt isolation:* Crypt, IEC, and ISC isolation, as well as organoid culture, were essentially done as described (Sato and Clevers, 2013). For adult crypt isolation duodenum tissue was rinsed with ice-cold PBS, cut open longitudinally, and villi were scraped off. Tissue was cut in ∼2 mm pieces and washed 5 times with ice-cold PBS. Tissue pieces were incubated in 2 mM EDTA in ice-cold PBS for 30-60 minutes. Crypts were isolated from up to 10 fractions after pipetting up and down 5 times with PBS. To isolate single cells for sorting, crypts were incubated in TrypLE (ThermoFisher) at 37 °C for 20-45 minutes. Single cells were stained and sorted, DAPI-negative and Epcam-positive cells were used for RNA-seq and ChIP-seq experiments. ISCs for clonal organoid outgrowth experiments were isolated by sorting DAPI-negative, GFP-high (top 5%) cells from *Lgr5-*EGFP mice.

*E18.5/P7/P21 IEC isolation:* Whole (E18.5) or proximal 10 cm (P7 and P21) small intestines were isolated and flushed with ice cold PBS when possible (P21). Small intestines were opened longitudinally (P21 and P7) and cut into small pieces that were washed with ice-cold PBS, and incubated with 2 mM EDTA in ice-cold PBS for 30 min. Whole epithelium was isolated by collecting all fractions, which was used directly for RNA isolation, for ChIP fractions were made single cell using TrypLE (ThermoFisher) and sorted as described. *Organoid cultures:* Organoids were grown and maintained in ‘basal crypt medium’ (Advanced DMEM/F12 medium supplemented with penicillin/streptomycin, 10 mM HEPES, 2 mM Glutamax, N2 (Thermo Fisher, 17502048), B-27 (Thermo Fisher, 17504044)) supplemented with N-acetyl-L-cysteine (Sigma, A7250), 50 ng/ml murine EGF [ThermoFisher, PMG8041], 20% R-spondin 1s conditioned medium (CM) (kind gift from Dr. Calvin Kuo), and 10% Noggin-CM (kind gift from Dr. Hans Clevers). For ISC clonal experiments, in the first 48 h after seeding, the medium was supplemented with Rock inhibitor (Y-27632) and Jagged-1 peptide (amino-acid sequence *CDDYYYGFGCNKFCRPR*, made in house, peptide synthesized as described (Bolscher et al., 2011)), 33% Wnt3-CM (kind gift from Dr. Hans Clevers) served as control. Medium was renewed every other day. For passaging, organoids cultures were washed, and matrigel and organoids were disrupted mechanically by strong pipetting, centrifuged at 200g, 5 min at 4 °C and resuspended in Matrigel to re-plate. Imaging of live organoids was done using an EVOS FL Auto 2. Structural Genomics Consortium supplied the inhibitors for the screen (www.thesgc.org), all of which are commercially available, concentrations are listed in Supplementary Table 1. Additionally, CHIR99021 (3 µM), IWP-2 (2 µM), VPA (1 mM) and DAPT (10 µM) were used.

*Vil-*Cre; *Yap*^f/f^*/Wwtr1*^f/f^ and *Vil-*CreERT2; *Yap*^f/f^;*Wwtr1*^f/f^ organoids were cultured in Basal Medium supplemented with NAC, B27, 50ng/ml human EGF (Peprotech, AF-100-15) and 100 ng/ml murine Noggin (Peprotech, 250-38) and either 500 ng/ml mouse RSPO1 (R&D systems, 3474-RS) or 10% RSPO1-conditioned medium. Established *Vil-*CreERT2; *Yap*^f/f^;*Wwtr1*^f/f^ organoids were cultured in the presence of 1 uM 4-OH-Tamoxifen (Sigma-Aldrich) for 72h prior to plating in the absence or presence of GSK-LSD1.

### Human organoids. Culture and staining

Human small intestine samples were obtained from patients undergoing elective surgery at Tokyo University Hospital with written informed consent. This was approved by the ethical committee (No. G3553-(7)). Crypt isolation and optimized organoid culture conditions that allow PC differentiation was done essentially as described^5,6^. In short, stroma was physically removed and the remaining epithelium was cut into 1-mm^3^ pieces, washed at least 5 times in ice cold PBS, and incubated in 2.5 mM EDTA in ice cold PBS for 1 hour. Isolated crypts were then suspended in Matrigel and seeded in 48-well plates. Domes of polymerized Matrigel were given the refined medium consisting of ‘basal crypt medium’ (see above) supplemented with 10 nM gastrin I (Sigma-Aldrich), 1 mM N-acetylcysteine (Sigma-Aldrich), 100 ng/ml recombinant mouse Noggin (PeproTech), 50 ng/ml recombinant mouse EGF (Thermo Fisher Scientific), 100 ng/ml recombinant human IGF-1 (BioLegend), 50 ng/ml recombinant human FGF-basic (FGF-2) (Peprotech), 1 mg/ml recombinant human R-spondin1 (R&D), 500 nM A83-01 (Tocris) and 50% Afamin-Wnt-3A serum-free conditioned medium. LGR5-iCaspase9-tdTomato organoids were made previously(Fujii et al., 2018). For staining, organoids were isolated from Matrigel using Cell Recovery Solution (Corning) and fixed in 4% paraformaldehyde for 20 min at room temperature. Next, organoids were washed with PBS, and permeabilized with 0.2% Triton X-100 in PBS for 20 min at room temperature. Blocking was done using Power Block Universal Blocking Reagent (BioGenex) for 20 min at room temperature, and rabbit anti-Lysozyme antibody (A0099, DAKO 1:1000 (Fig. 1h), GTX72913, GeneTex, 1:200 (Fig. 2f)) and anti-RFP (600-401-379, Rockland 1:500) was incubated overnight at 4 °C. Organoids were washed 3 times with PBS, and secondary antibody incubation was done for 30 min at room temperature. Nuclear counterstaining was done simultaneously with secondary antibody incubation using Hoechst 33342 (Thermo Fisher Scientific). Stained organoids were suspended in 1 drop of ProLong Diamond Antifade Mountant (Thermo Fisher Scientific) and mounted onto a 35-mm glass bottom dish. Images were captured using a confocal microscope (SP8, Leica).

### Immunohistochemical staining of intestinal tissue

For immunohistochemical staining and imaging, tissues were harvested and fixed in swiss rolls. After fixation in formalin, tissues were embedded in paraffin and cut in 4 µm sections. Paraffin sections were rehydrated and peroxidase activity was blocked in 3% hydrogen peroxide. Antigen retrieval was performed in citrate buffer pH6. Sections were stained overnight with primary antibodies against Ki67 (1:500, Thermo Scientific MA5-14520), Lysozyme (1:750, Dako A0099), Sox9 (1:200, Millipore), LSD1 (1:200, Cell Signalling 2184S), H3K4me1 (1:100, Cell Signaling 9723) and H3K4me2 (1:1500, Cell Signalling 9725). The sections were washed in TBS and Tween-20 and stained for 1 hour with HRP-labelled secondary antibody (Dako K4003). The staining was developed with diaminobenzidine (DAB) chromogenic substrate (Dako K5007) and mounted with Glycergel mounting medium (Dako C056330). Tissues were imaged using a Nikon eclips Ci-L microscope.

### Immunofluorescence staining of intestinal tissue and organoids

For immunofluorescence labeling and imaging, tissues (first 5 cm of duodenum) were harvested and fixed in swiss rolls. After fixation in formalin, tissues were paraffin embedded and cut in 4 µm sections. Briefly, paraffin sections were treated as before for IHC, and after antigen retrieval were blocked and permeabilized in PBS with TX-100 0.2%, Normal Goat Serum (NGS) 2%, BSA 1% and Tween-20 0.05%. Sections were then stained overnight in the same blocking buffer with primary antibodies against GFP (1:2000 Abcam 13970), YAP (1:200 Cell Signaling Technologies 14074S), OLFM4 (1:200 Cell Signalling 39141S) or LSD1 (1:200 Cell Signaling Technologies 2184S). Tissues were then incubated with the corresponding secondary antibodies for 3 hours (1:500 Alexa Fluor), Rhodamine-labeled UEA1 (5 µg/ml Thermo Fisher Scientific NC9290135) and Hoechst 33342 (1:10,000). Washes were performed with PBS + Tween-20 0.1%.

For organoid staining, organoids were grown in Matrigel on eight-chamber μ-slides (Ibidi 80826) and fixed after exposition to the specific treatments in PBS containing 4% paraformaldehyde (pH 7.4) and 2% sucrose for 20-30 min, permeabilized (PBS, 0.2% Triton X-100) and blocked (PBS-Triton X-100 0.2%, 2% NGS, 1% BSA). Primary antibodies against the following antigens were used, diluted in the same blocking buffer: Lysozyme (1:500, Dako A0099), GFP (1:2000 Abcam 13970) and LSD1 (1:400 Cell Signaling Technologies 2184S) overnight at 4°C with slow agitation. Rhodamine-labeled UEA1 (5 µg/ml Thermo Fisher Scientific NC9290135) and Hoechst 33342 (1:10,000) were used to stain secretory cells and nuclei respectively together with the corresponding secondary antibodies (1:500 Alexa Fluor) and incubated overnight in PBS with 0.2% Triton X-100, 1% NGS and 0.5% BSA at 4°C. Tissue sections and organoids were both mounted using Fluoromount G (ThermoFisher Scientific, 00-4958-02) and imaged with a Zeiss 510 Meta Live or a Zeiss LSM880 confocal microscope, using 20x and 40x objective lens.

### In situ hybridization

In situ hybridization was performed on FFPE tissues using RNAscope® 2.5 HD BROWN reagent kit (Advanced Cell Diagnostics (ACD) 322371). Tissue sections (4µm) were deparaffinised with Neoclear and 100% ethanol. The slides were pretreated with hydrogen peroxide for 10 minutes, target antigen retrieval reagent for 15 minutes, and protease plus reagent for 30 minutes (ACD 322300 and 322000). The sections were hybridized with probes for Mm-Wnt3 (ACD 312241), Mm-Olfm4 (ACD 311831), Mm-Atoh1 (ACD 408791),m positive control Mm-Ppib (ACD 313911) and negative control Mm-DapB (ACD 310043). For amplification and chromogenic detection the 2.5 HD Detection Reagents BROWN kit (ACD 322310) was used. The slides were counterstained with hematoxylin, dehydrated and mounted with Neomount (Merck 109016). Tissues were imaged using a Nikon eclips Ci-L microscope.

### Flow cytometry analysis of organoids

Organoids were mechanically disrupted, centrifuged at 2000 rpm and incubated with TripLE (ThermoFisher) at 37 degrees Celsius for 50 minutes for dissociation into single cells. Cells were incubated with DAPI and analyzed using a flow cytometer (FACSCanto II; BD). Stem cell populations were gated as DAPI negative and GFP-high (top 5%) and analyzed using FlowJo software.

### Western Blot

Organoids were harvested in lysis buffer (1% NP-40, 0.02% SDS in 1X TBS) on ice for 30 minutes. Debris was pelleted by spinning down at 14 000 RPM for 30 min. Supernatant was diluted in 4x NuPage sample buffer with 100 mM DTT, and samples were run using precast 4-12% gels using the NuPage system, and blotted using iblot 2 (all ThermoFisher). Membranes were incubated with antibodies against Lysozyme (Dako A0099), Tubulin (Abcam Ab6046). Secondary antibodies (HRP linked) were swine anti–rabbit (P039901-2) and goat anti–mouse (P044701-2) (DAKO). Imaging was done using SuperSignal West Femto (ThermoFisher) on a Lycor machine. Bands were quantified using Image Studio software.

### qPCR

RNA from organoids was isolated using either an RNeasy kit (Qiagen) or Quick-RNA kit (Zymo). Reverse transcription was carried out by using the High-Capacity RNA-to-cDNA Kit (ThermoFisher). Quantitive PCR was performed using the QuantiFast SYBR Green PCR Kit (Qiagen) using primers for *Hprt* (fwd: cctcctcagaccgcttttt, rev: aacctggttcatcatcgctaa), *Actb* (fwd: actaatggcaacgagcggttc, rev: ggatgccagaggattccatacc), *Lyz1* (fwd: ggcaaaaccccaagatctaa, rev: tctctcaccaccctctttgc), *Lyz1* (fwd: gccaaggtctacaatcgttgtgagttg, rev: cagtcagccagcttgacaccacg), Defa, fwd: aatcctcctctctgccctcg, rev: accagatctctcaatgattcctct), *Yap1* (fwd: tggccaagacatcttctggt, rev: caggaacgttcagttgcgaa), *Wwtr1* (fwd: tggggttagggtgctacagt, rev: ggattgacggtcatgggtgt), *Gapdh* (fwd: tgttcctacccccaatgtgt, rev: tgtgagggagatgctcagtg), *Olfm4* (fwd: ggatcctgaacttttggtgct, rev: acgccaccatgactacagc), and *Wnt3* (fwd: ctcgctggctacccaattt, rev: gaggccagagatgtgtactgc). Samples were commonly analyzed in duplicate and RNA expression was calculated either normalized to reference gene, or additionally normalized to control conditions.

### RNAseq preparation

RNA for WT and LSD1KO crypts was isolated by sorting IECs (DAPI-, Epcam+) in 2x RNA shield buffer (Zymo) and RNA isolation using the Quick-RNA Micro prep kit (Zymo). Library preparation was done using the Illumina TruSeq Stranded protocol, and samples were sequenced at 75×2 bp PE reads on an Illumina NS500 MO flow-cell. Sequencing was performed by the Genomics core facility (GCF, NTNU). Crypts from E18.5, P7 and E18.5 was directly dissolved in RNA isolation buffer and RNA was isolated using the Quick-RNA micro prep kit (Zymo). Library preparation was done using the NEB Next® Ultra™ RNA Library Prep Kit. Sequencing was performed by Novogene (UK) Co.

### RNAseq analysis

Sequenced reads were aligned with STAR to the *Mus musculus* genome build mm10 (Dobin et al., 2013; Frankish et al., 2019). The number of reads that uniquely aligned to the exon region of each gene in GENCODE annotation M18 of the mouse genome was then counted using featureCounts (Liao et al., 2014). Genes that had a total count less than 10 were filtered out. Differential expression was then determined with DESeq2 using default settings (Love et al., 2014). Interesting differential genes were plotted with a volcano plot using the R package EnhancedVolcano. Heatmaps where generated using the R-package pheatmap. Count values for each gene were transformed to rates per bp by dividing the count for a gene by the length of the total exon region for that gene. Rates per bp where then converted to Transcripts Per Million (TPM) by dividing the rate per bp for each gene by the sum of rates per bp for all the genes in that sample and multiplying with one million. PCA analysis was performed using the function sklearn.decomposition.PCA in scikit-learn. Gene set enrichment analysis (GSEA) was done by sorting the output from DESeq2 by log2 fold change and with the log2 fold change as weights. GSEA was run with the R package clusterProfiler using 10000 permutations and otherwise default settings. Gene sets were generated from published datasets and can be found in supplementary table 3.

### Microarray analysis

Gene expression in human small intestinal organoids was analyzed using the PrimeView Human Gene Expression Array. Raw expression data were normalized with the rma function in the R/Bioconductor package affy and the normalized values were used to calculate log fold change(Gautier et al., 2004). For each gene, the probe with the highest absolute log fold change was used. GSEA was run on this list of genes as described for the RNAseq analysis.

### ChIP-seq

DAPI-negative, Epcam-positive IECs were sorted in PBS containing 20 mM Sodium Butyrate, and cross-linked by incubation in 1% formaldehyde for 8 minutes. Glycine was added to a final concentration of 125 mM and incubated for 5 minutes at room temperature. Using a swing-out rotor cells were washed 3 times in ice-cold PBS with 20 mM Sodium Butyrate. After washing, cells were snap frozen in liquid nitrogen and stored in −80 °C. The ChIP-seq was carried out similarly to previously described protocols (Dahl and Collas, 2008) *Binding of antibodies to paramagnetic beads.* The stock of paramagnetic Dynabeads Protein A was vortexed thoroughly to ensure a homogenous suspension before pipetting. Dynabeads stock solution (5 μL per IP) was transferred into a 1.5-ml tube, which was placed on a magnetic rack and the beads captured on the tube wall. The buffer was discarded, the beads washed twice with 200 μL standard RIPA buffer (10 mM Tris-HCl pH 8.0, 140 mM NaCl, 1 mM EDTA, 0.5 mM EGTA, 1% Triton X-100, 0.1% SDS, 0.1% Na-deoxycholate) and resuspended in standard RIPA buffer to a final volume of 100 uL per IP. 99 μL of this was aliquoted into each 0.6 mL tube on ice, and antibody (1.2 ug of anti-H3K4me1: Diagenode, C15410194, Lot A1862D or 4 uL anti-H3K4me2: Cell Signaling #9725, Lot 9) was added per 0.6 mL-tube. Tubes were then incubated at 40 r.p.m. on a ‘head-over-tail’ tube rotator for at least 16h at 4 °C.

#### Chromatin preparation, Lsd1 cre+ / cre-

Crosslinked cell pellets containing 335 000-500 000 cells were thawed on ice. The 6-10μL pellets were added Lysis buffer (50 mM Tris– HCl pH 8.0, 10 mM EDTA pH 8.0, 1% SDS, 20 mM sodium butyrate, 1 mM PMSF and protease inhibitor cocktail) to a total of 160 μL and incubated on ice for 10 min. The samples were sonicated for 8 x 30 s using a UP100H Ultrasonic Processor (Hielscher) fitted with a 2-mm probe. We allowed for 30 s pauses on ice between each 30 s session, using pulse settings with 0.5 s cycles and 27% power. After the final sonication, 340 μL standard RIPA (with 20 mM sodium butyrate, 1 mM PMSF and protease inhibitor cocktail) was added to the tube while washing the probe, followed by thorough mixing by pipetting. 20 μL was removed as input, and the remaining solution was diluted further with 1mL standard RIPA buffer (with 20 mM sodium butyrate, 1 mM PMSF and protease inhibitor cocktail). The samples and inputs were centrifuged at 12,000 g in a swinging-bucket rotor for 10 min at 4 °C and the supernatants were transferred to a 1.5-ml tube on ice. 66 000 – 100 000 cells were used per IP.

#### Immunoprecipitation and washes

Pre-incubated antibody–bead complexes were washed twice in 200 μl standard RIPA buffer by vortexing roughly. The tubes were centrifuged in a mini-centrifuge to bring down any solution trapped in the lid and antibody–bead complexes were captured in a magnetic rack. After removal of RIPA, 177-500 μl of chromatin (equivalent of 50,000-100,000 cells per ChIP) was added to each tube, then incubated at 4 °C, 40 r.p.m. on a ‘head-over-tail’ rotator for at least 16 h. For H3K4me1 ChIPs, the chromatin– antibody–bead complexes were washed three times in 100 μl ice-cold standard RIPA buffer. For H3K4me2 ChIPs, the reactions were washed once in standard RIPA, once in RIPA with increased salt and SDS (300 mM NaCl and 0.20% SDS), once in RIPA with increased salt and SDS (300 mM NaCl and 0.23% SDS), once in standard RIPA. All washing steps were performed with 100uL ice cold buffer supplemented with 20 mM sodium butyrate, 1 mM PMSF and protease inhibitor cocktail. Each wash involved rough vortexing at full speed, repeated twice with pauses on ice in between. Next, a wash in 100μl TE and tube shift was carried out.

#### DNA isolation and purification

TE was removed and 150μl ChIP elution buffer was added (20 mM Tris-HCl pH 7.5, 50 mM NaCl, 5 mM EDTA. 1% SDS, 30 μg RNase A) and incubated at 37 °C, 1 h at 1,200 r.p.m. on a Thermomixer. The input samples were added ChIP elution buffer up to 150uL and incubated similarly. 1μl of Proteinase K (20 mg/ml stock) was added to each ChIP or input tube and incubated at 68 °C, 4 h at 1,250 r.p.m. The ChIP eluates were transferred to a 1.5-ml tube. Then, a second elution with 150μl was performed for 5 min and pooled with the first supernatant. The ChIP and input DNA was purified by phenol-chloroform isoamylalcohol extraction, ethanol-precipitated with 11μl acrylamide carrier and dissolved in 10-15μl EB (10 mM Tris-HCl).

#### Library preparation and sequencing

ChIP and input library preparations were performed according to the QIAseq Ultralow Input Library Kit procedure. Sequencing procedures were carried out according to Illumina protocols, on a NextSeq 500 instrument, with 75bp single end reads using high output reagents. The sequencing service was provided by the Norwegian Sequencing Centre (www.sequencing.uio.no).

### ChIP-seq analysis

Sequence reads were deduplicated with BBMaps clumpify tool and then aligned with STAR to the *Mus musculus* genome build mm10 (Dobin et al., 2013; Frankish et al., 2019) (*Bushnell, B. BBMap. SourceForge Available at: https://sourceforge.net/projects/bbmap Accessed: 12th February 2019*). Peaks were identified using Model-Based analysis of ChIP-seq 2 (MACS2) with peak type set to broad and genome size 2652783500 (Zhang et al., 2008). Input files where supplied for H3K4me1/2. Peaks from all samples that were compared in differential expression were merged with BEDTools to create a union set and featureCounts was used to count the number of reads, including multi-mappers, for each sample in the union set of peaks (Zhang et al., 2008). Differential peaks was determined from the counts with DESeq2 (Love et al., 2014) using default settings and. deepTools2 was used to create heatmaps (Ramírez et al., 2016). Peak locations were associated with the gene that has the closest transcriptional start site (TSS) with the closest command in bedtools (Quinlan and Hall, 2010). Ties were resolved by only reporting the first hit. TSS sites were downloaded from biomart for GRCm38.p6. The peaks were grouped on the distance to the TSS and the size of each group was plotted. The list of differential peaks with associated genes was grouped by gene and sorted on the differential peak that had the smallest p-value for each gene. Each gene was determined to be either up or down in signal based on whether the total change was above or below zero, where total change is defined as the average of log fold change multiplied by peak length of all peaks associated with that gene. Venn diagrams where created with the R package Euler (Larsson2018). Bigwig files describing the score across the genome were created with deepTools2 and scaled to the count of the sample with the least aligned reads for each group (e.g. H3K4me1, H3K4me2) (Ramírez et al., 2016). Heatmaps of regions of interest was created with deepTools2. Chipseq profiles were created in integrative genomics viewer (IGV) (Robinson et al., 2011).

### Fetal RNAseq (from Yui et al)

Published microarray raw data was downloaded from ArrayExpress under the accession number “E-MTAB-5246”, normalized with “neqc” function in the R package limma and then log2fc was calculated from the normalized expression values (Yui et al., 2018; Ritchie et al., 2015). GSEA was performed as described in the RNAseq methods.

### scRNAseq analysis

Preprocessed and normalized scRNAseq data where downloaded from GSE117783 (Ayyaz et al., 2019). The control treated cells where randomly sub sampled so the two groups had equal number of cells and the density of LSD1 expressing cells where plotted in base R. Y-axis is shortened to show distribution of cells that has an expression larger than zero.

### Statistical analysis

Statistical significance was determined either using Student’s *t* test or 1-way ANOVA with Tukey’s post hoc test, or, when n<10 non-parametric testing (Mann Whitney test) was done. Significance levels are indicated in figure legends.

## Data availability statement

All raw data is available through ArrayExpress. WT/KO Crypt RNAseq: E-MTAB-7862, WT/KO Crypt ChIP seq: E-MTAB-7871, E18/P7/P21 RNA-seq E-MTAB-8713, and Human microarray: E-MTAB-7871.

